# Fate-mapping lymphocyte clones and their progenies from induced antigen-signals identifies temporospatial behaviours of T cells mediating tolerance

**DOI:** 10.1101/2023.03.16.532070

**Authors:** Munetomo Takahashi, Tsz Y. So, Kate Williamson, Zhaleh Hosseini, Katarzyna Kania, Michelle Ruhle, Tiffeney Mann, Martijn J. Schujis, Paul Coupland, Dean Naisbitt, Timotheus Y.F. Halim, Paul A. Lyons, Pietro Lio, Klaus Okkenhaug, David J. Adams, Ken G.C. Smith, Duncan I. Jodrell, Michael A. Chapman, James E. D. Thaventhiran

## Abstract

Tissue homeostasis is maintained by the behaviours of lymphocyte clones responding to antigenic triggers in the face of pathogen, environmental, and developmental challenges. Current methodologies for tracking the behaviour of specific lymphocytes identify clones of a defined antigen-receptor—antigen binding affinity. However, lymphocytes can receive antigenic signals from undefined or endogenous antigens, and the strength of each signal, even for the same lymphocyte, varies with accessory signalling, across tissues and across time. We present a novel fate-mapping mouse, that, by tracking lymphocyte clones and their progenies from induced antigen signals, overcomes these hurdles and provides novel insights into the maintenance of tissue homeostasis. We demonstrate the systems use by investigating the maintenance of localised T cell tolerance in tumour immunity. In a murine tumour model, our system reveals how Tregs differentiate to a reversible, tolerance inducing state within the tumour, and recirculate, while CD8+ T cells failing to recirculate, differentiate to an increasingly exhausted, tolerant state in the tumour. These contrasting T cell behaviours provide means by which immunity can tolerate a particular anatomical niche while maintaining systemic clonal protection. Our system can thus explore lymphocyte behaviours that cannot be tracked by previous methods and will therefore provide novel insights into the fundamental mechanisms underlying immunity’s role in tissue homeostasis.

## Introduction

Over the past two decades, there has been an increasing appreciation for the central role of adaptive immunity in regulating tissue homeostasis in non-infectious diseases. While it is clear that lymphocyte responses are fundamental to autoimmune diseases, recent research has substantiated a more central role for adaptive immunity in conditions such as cancer(*1*), cardiovascular disease(*2*), metabolic disorders(*3*), dementia(*4*) and even psychiatric illness(*5*, *6*). In contrast to infectious diseases settings, lymphocytes in these settings are reactive to undefined or endogenous antigens present in the host, leading to difficulty tracking their responses. While significant investments in therapeutic targeting of adaptive immunity have led to treatments for previously incurable diseases(*7*), the lack of success in other disease(*8*, *9*) suggest there is substantial room for improvement. A deeper understanding of the immune response to antigens in non-infectious disease settings will prove crucial in further broadening the success of immunotherapy.

Adaptive immunity is understood according to the precepts of the clonal selection theory, which proposed that lymphocyte clones are selected by antigen to proliferate, differentiate, and persist to mediate protection directed against antigen-bearing pathogens(*10*). However, at the time of this proposal, recognition by the antigen-receptor was thought the sole triggering event capable of distinguishing pathogenic, exogenous, microbial derived ‘non-self’ from ‘self’ antigens. Since then, our understanding of clonal selection has been substantially revised(*11*, *12*). We now understand that all lymphocyte clones undergo positive selection via their antigen receptor for ‘self’ reactivity and can only persist with on-going self-reactivity(*13*). Inflammatory lymphocyte clones with autoimmune potential therefore exist within the normal repertoire(*14*, *15*), but in health, the quiescence of these clones relies on the balance of positive and negative accessory signals. Critical positive accessory signals required for clone selection are delivered by costimulatory signalling that can increase the signal from self-or foreign-antigen through the B and T cell antigen-receptor(*16*–*18*). Countering this, signalling through the antigen-receptor is also limited, either directly(*19*) or indirectly(*20*) by immune checkpoints. Inflammatory lymphocyte responses are also held in check by the negative signals of clonally expanding T regulatory (Treg) cells which respond to-self(*21*) and non-self-antigens(*22*).Therefore, for any given antigen, the signal received by the lymphocyte via its antigen receptor varies with this accessory signalling and can therefore vary in different tissues and at different times.

The development of the first TCR/BCR transgenic lymphocytes(*23*–*25*) and then tetramers(*26*, *27*) permitted the study of clonal selection by tracking the behaviour of reactive lymphocytes challenged with cognate antigen. Whilst most of what we know about adaptive immune responses has been learnt from these systems, they have limitations; TCR/BCR-transgenics generally track the behaviour of a single clone in response to a high-avidity antigenic trigger, whilst tetramers track the behaviour of a polyclonal response, the avidity threshold required for tetramer binding is higher than that required for T cell activation(*28*, *29*). Therefore, these systems, although able to track the antigen-selection of clones responding to an inoculated high-avidity exogenous antigen found in infectious disease, have difficulties in tracking the lymphocyte clones selected by antigen at different times or sites during a chronic immune response to persisting undefined or endogenous antigen, found in conditions such as cancer and autoimmunity.

A potential strategy to overcome these limitations is to track antigen-receptor signalling based on the actual signal received by the lymphocyte, rather than relying on the avidity of the antigen receptor. This can be achieved by fate-mapping antigen-receptor signalling itself, which has previously been visualized using fluorescent reporters of the Nr4a1 (Nur77) gene. This gene’s promoter activity is quantitatively driven by the strength of antigen-receptor ligation in B and T lymphocytes(*14*, *30*). Using these mice, Zikherman et al., visualised the self-reactivity and restraint of autoreactive B lymphocytes at steady state. However, these mice could only report current antigen-receptor ligation and had no ability to track the clonal progeny driven by antigen-selection.

To visualise and understand clonal selection, we have developed and validated the antigen-receptor signalling reporter mouse (AgRSR). This novel transgenic mouse links signalling from the antigen-receptor of lymphocytes to inheritable lymphocyte clone marking, allowing direct visualization of the clonal selection theory *in vivo*. We have validated the AgRSR mouse by demonstrating its ability to fatemap antigen signalled T cells and their progenies. We applied the system to a murine tumour model, to show how immune system can maintain tolerance within a tumour, while preserving its capacity to protect against metastatic dissemination. Through this demonstration, we propose that the AgRSR mouse system will provide novel insights into diverse diseases in which adaptive immunity plays a significant role.

## Results

In the AgRSR mouse, the promoter of a T and B cell receptor response gene, Nur77 (*NR4A1*) drives equimolar expression of a red fluorescent protein (Katushka), and Cre-ERT2 recombinase. We first assessed the requirements for fluorescence in T cells. We crossed the AgRSR strain to the OVA specific OT-I TCR transgenic strain, for which variant peptide ligands of the TCR have been characterised(*13*). Naïve CD8+ T cells from these animals were stimulated with OVA peptide variants *in vitro*. Consistent with results from a previous Nur77-GFP(*30*) strain, the level of Katushka induced by each ligand directly correlated with its stimulatory activity (Figures 1A and 1B), indicating Katushka expression is dependent on TCR signalling. This suggested that the AgRSR mouse could fate-map T cells that had received TCR signalling.

**Figure 1.**
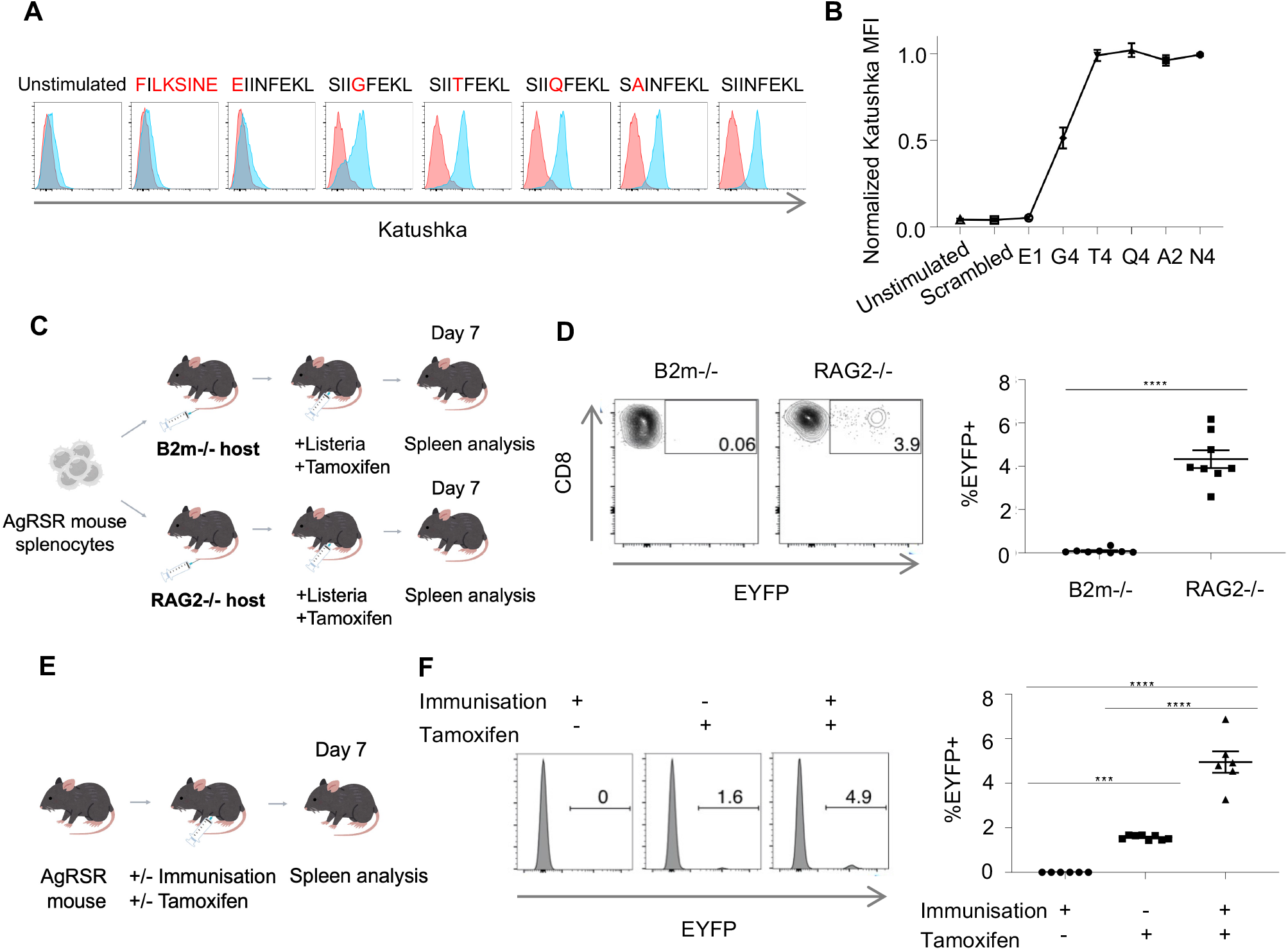
The AgRSR mouse fate-maps T cells based on antigen-signaling. (A-B) Range of Katushka expression of OT-I AgRSR and OT-I WT CD8+ T cells, following stimulation with the indicated peptides, as displayed by flow cytometry histograms (A) and normalised fluorescence (B). Data are representative of at least two independent experiments. (C) Schematic of experimental design; AgRSR-LSL-EYFP splenocytes were adoptively transferred to CD8+ T cell deficient B2m-/- and RAG2-/- strains, challenged with Listeria and tamoxifen treated. (D) Representative flow cytometry plots of EYFP and CD8 expression of splenocytes taken from the indicated recipient mouse strain at day 7 post infection (Left), and summary plot of all experiments (Right). Data are pooled from two independent experiments (n=7 per condition). (E) Schematic of experimental design; AgRSR-LSL-EYFP mice were treated with/without immunisation and with/without tamoxifen. (F) Representative flow cytometry histograms of EYFP expression in CD8+ T cells taken from splenocytes from the indicated mice at day 7 post immunisation (Left) and summary plot of all experiments (Right). Data are pooled from three independent experiments (n=6 per condition). Dots represent a single mouse (D, F). Mean ± SEM as shown (B, D, F). Statistical testing via unpaired two-tailed students t-test (****, p <0.0001; ***, p <0.001; **, p <0.01; *, p<0.05; ns, p>0.05).

Crossing AgRSR mice to Rosa-Lox-Stop-Lox (LSL)-EYFP strains generates AgRSR-LSL-EYFP mice in which TCR signalling, coupled with tamoxifen administration, permanently marks T cells and their progenies. Nurr77 expression dependent Cre-ERT2, in the presence of tamoxifen, excises a stop cassette that otherwise prevents EYFP expression from a constitutively active promoter. Since this is a DNA recombination event, the expression of EYFP is inherited in all progenies. We transferred splenocytes from AgRSR-LSL-EYFP mice into B2m knockout (KO) and RAG2 KO host mice to assess the TCR signalling dependence on functional Cre activity (Figure 1C). In these CD8+ T cell deficient strains, the MHC-I TCR-ligand is absent in the B2m but present in the RAG2 KO mice. Recipient mice infected with Listeria and treated with tamoxifen showed EYFP fluorescence in CD8+ T cells from the RAG2 KO but not the B2m KO mice (Figure 1D), demonstrating specificity of recombination to TCR signalling. We next confirmed dependence of recombination on tamoxifen administration by observing a significant percentage of CD8+ T cells EYFP+ in tamoxifen-treated mice post-immunisation and no detectable fluorescence in its absence (Figures 1E and 1F). Taken together, these results validate the use of the AgRSR mouse in fate-mapping antigen selected T cells.

The post-expansion behaviours of T cell clones establish the confines of reactivity and tolerance, and the AgRSR system enables the tracking of these behaviours following antigen-ligation at specific times, and, by localised administration of tamoxifen, specific tissues. Tumour tolerance forms after tumour establishment, suggesting the behaviour of T cell clones receiving antigen signals after tumour establishment play a major role in its maintenance. We therefore utilised the AgRSR mouse to investigate the spatiotemporal behaviours of clones responding to these antigens.

The YUMMER1.7 melanoma model provides a persistent immunological challenge that has been used for the characterization of CD8+ T cell responses during immunotherapy(*31*–*33*). When implanted in tamoxifen treated AgRSR mice, tumours grew consistently (Figure S1A). Only AgRSR mice receiving tamoxifen after, and not before tumours became palpable showed elevated CD8+ T cell tumour EYFP+ frequency (Figures 2A and 2B). In these mice, splenic EYFP+ CD8+ T cells expressed PD-1 and tumour EYFP+ CD8+ T cells were PD-1Hi (Figures 2C and 2D) indicating substantial EYFP enrichment of T cell clones responding to tumour antigens in both populations(28, 29). We quantified the longitudinal changes in EYFP+ frequency of CD8+ and CD4+ T cells in the secondary lymphoid system and tumour (Figures 2E, 2F and S1B). After TCR activation in the draining lymph node, antigen-selected T cell clones expand, and their cells recirculate through the lymphatics and blood stream prior to entry into the tumour. EYFP+ frequencies in all compartments followed similar expansion kinetics, indicating systemic labelling of antigen-signalled T cells across different tissues. From day 8 post-tamoxifen, EYFP+ CD8+ T cell frequency plateaued as clones ended their initial expansion and entered a state of persistence.

**Figure 2.**
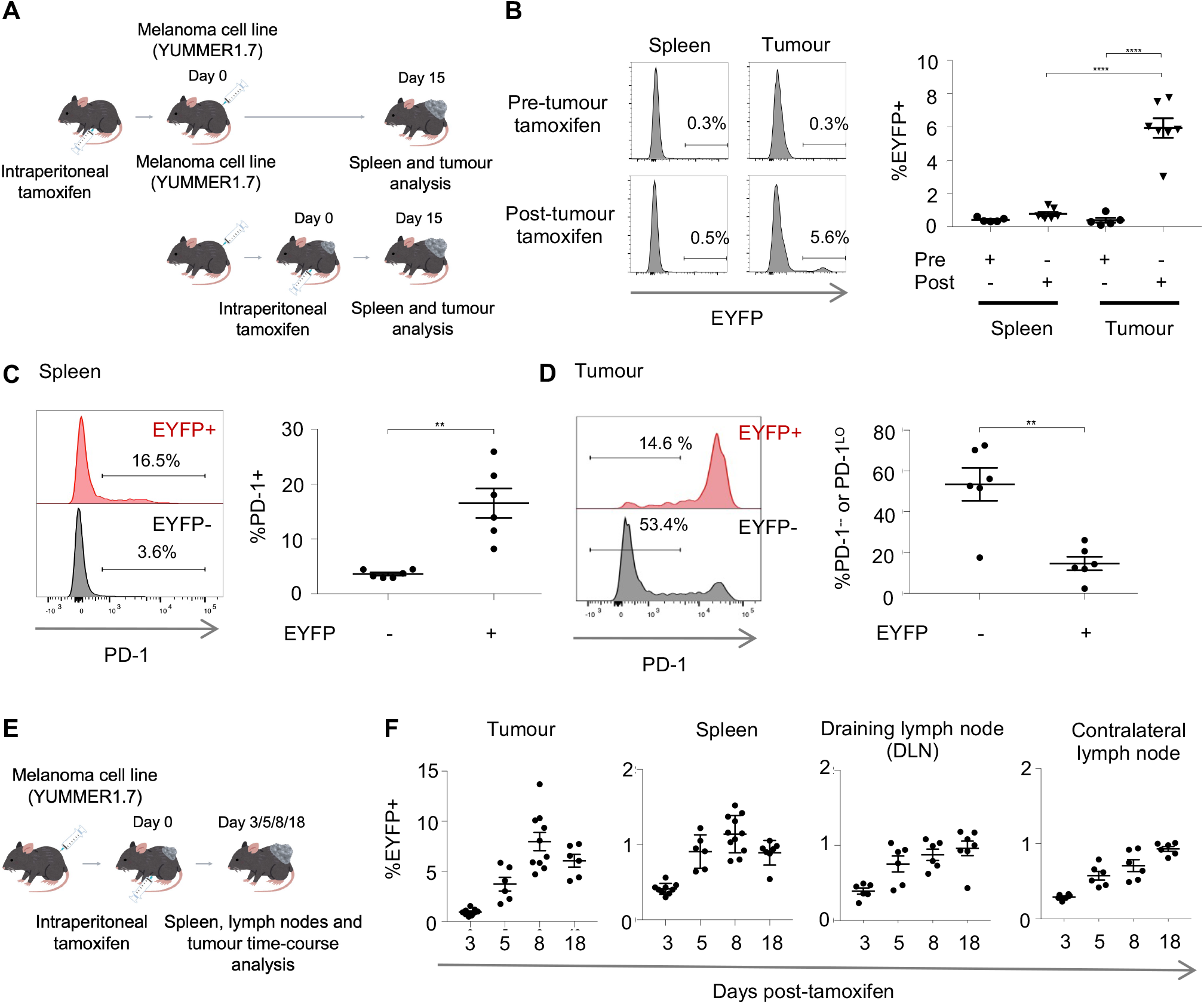
The AgRSR mouse tracks antigen signalled T cells in the tumour immune response. (A) Schematic of experimental design; AgRSR-LSL-EYFP mice were tamoxifen treated either before (pre-tumour) or after (post-tumour) subcutaneous injection of Yummer 1.7 melanoma tumour cells, and CD8+ T cells were assessed 15 days afterwards. (B) Representative flow cytometry histograms (Left) and summary plot of all experiments (Right). Data are pooled from two independent experiments (n=? per condition). (C-D) Representative flow cytometry histograms (Left) and summary plot (Right) of PD-1 expression in CD8+ T cells from the spleen (C) and the tumour (D) of EYFP+ and EYFP-CD8+ T cells 8 days after tamoxifen labelling in tumour bearing mice. Gated to show the proportion of CD8+ T cells expressing PD-1 in the spleen (C) and lacking high PD-1 expression in the tumour (D). Data are pooled from two independent experiments (n=6). (E) Schematic of experimental design; AgRSR-LSL-EYFP mice were implanted with Yummer 1.7 melanoma tumour cells, treated with tamoxifen at day 7, and CD8+ T cells in the spleen, tumour, ipsilateral inguinal lymph node (draining lymph node (DLN)) and contralateral lymph node (non-draining lymph node (nDLN)) were sampled at days 3, 5, 8 and 18. (F) Results of the temporal assessment of EYFP+ percentages of CD8+ T cells in the indicated tissues. Data are pooled from four independent experiments (n≥5 per condition). Dots represent a single mouse (B, C, D, F). Mean ± SEM as shown (B, C, D, F). Statistical testing via paired two-tailed students t-test and ordinary one-way ANOVA (****, p <0.0001; ***, p <0.001; **, p <0.01; *, p<0.05; ns, p>0.05).

To investigate the behaviours of expanded clones in both the effector site and the circulating immune system, we analysed EYFP+ T cells from the tumour and spleen of 4 mice by paired scRNA-seq and T-cell receptor (TCR) sequencing, 8 days after tamoxifen injection (Figure 3A). Naïve T cells carry a unique TCR sequence which is inherited by its progenies during clonal expansion. The TCR sequence therefore provides a genetic barcode, which, in conjunction with EYFP labelling, enables the tracking of individual clonal expansions of T cells after antigen signalling. In the dataset, most cells from the spleen were nonexpanded whereas most cells from the tumour were members of expanded clonal populations (Figure S2A). Non-expanded cells highly expressed markers associated with a naïve state suggesting these cells had received sufficient antigen signals to recombine, but insufficient coreceptor signals to clonally expand or differentiate. We focused our analysis on expanded clonal populations (Figure 3B). Filtering the dataset for the largest (>15 cells) clonal populations enabled us to distinguish CD8+ and CD4+ T cell clonal populations by their Cd8a and Cd4 expression (Figures 3C, 3D and S2B). Based on the proportion of cells expressing Foxp3 (Figures 3E and S2C), we could further categorise CD4+ T cell clonal populations as populations that did not express Foxp3, populations that transiently expressed Foxp3 (<40% of the cells expressing Foxp3), and populations that persistently expressed Foxp3 (>40% of the cells expressing Foxp3). Populations persistently expressing Foxp3 were significantly less expanded in the spleen and more expanded in the tumour (Figure S2D) but all populations were equally distributed across all mice (Figure S2E). Compared to Foxp3+ cells from populations that transiently expressed Foxp3, Foxp3+ cells from populations that persistently expressed Foxp3 upregulated genes associated with known effector Treg states (including Helios, Gzmb, Ctla4) (Figures 3F and S2F). These Foxp3+ cells also made-up most of the Foxp3+ cells in the tumour (Figure S2G). These results indicate that in addition to marking CD8+ T cell clonal populations, the AgRSR enables the tracking of both transient and stable clonally expanded Treg clonal populations.

**Figure 3.**
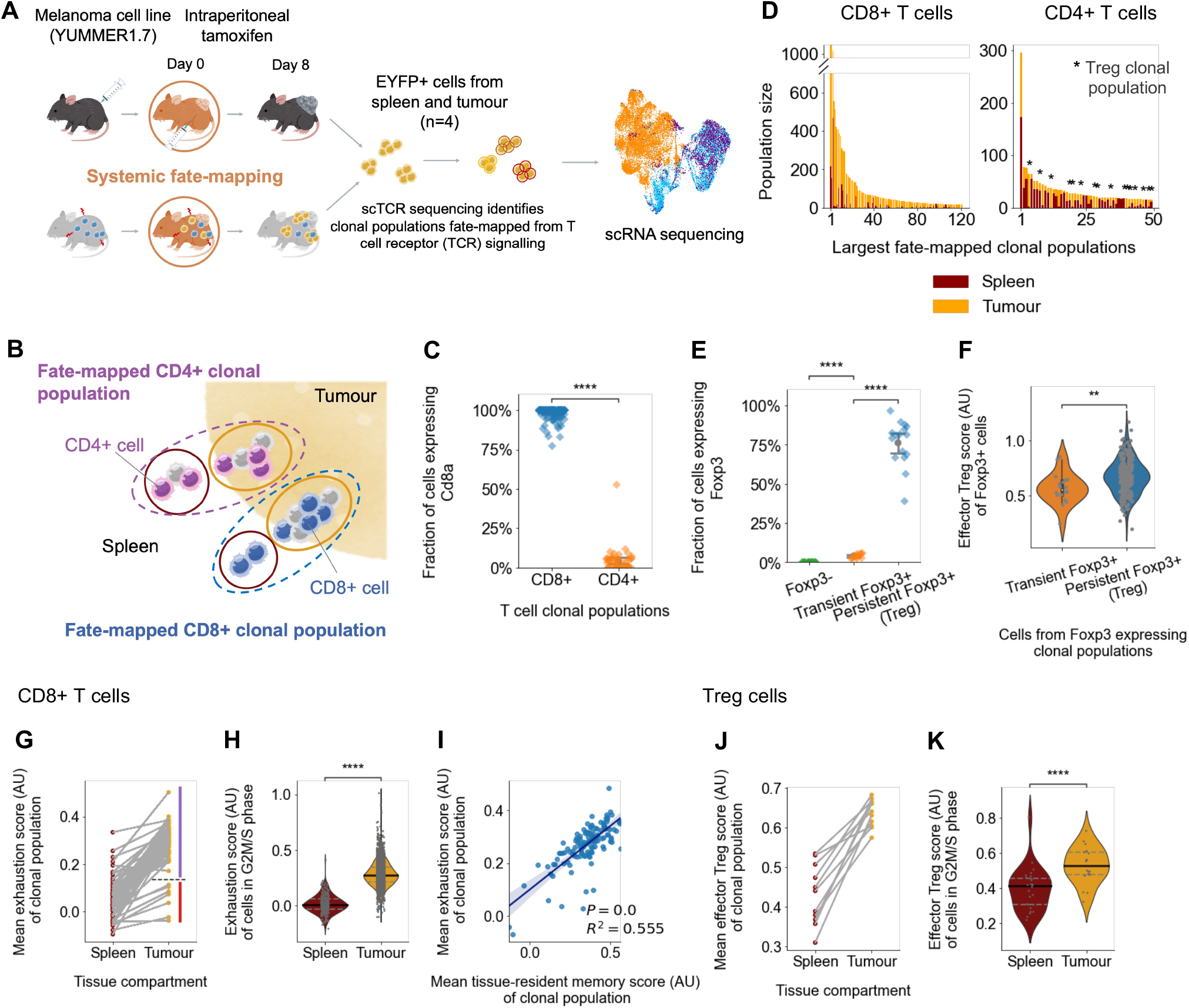
CD8+ T and Treg cell states mediating tolerance are partitioned on tissue site. (A) Schematic of experimental design; 8 days after intraperitoneal tamoxifen injection EYFP+ T cells were sorted from the spleen and tumour of AgRSR-LSL-EYFP mice and subject to single-cell RNA and VDJ analysis (n=4). (B) Schematic displaying captured T cells clonal populations being grouped into CD4+ and CD8+ T cell clonal populations based on CD4 and CD8a expression. (C) Fraction of cells expressing Cd8a for the largest (population size greater than 15) CD8+ and CD4+ T cell clonal populations. Clonal populations express Cd8a if at least one cell is Cd8a+. (D) Largest fate-mapped CD8+ and CD4+ T cell clonal populations ranked by size and coloured by the numbers of cells within the tumour or spleen compartments. (E-F) Foxp3+ expression analysis reveals three categories of CD4+ T cell clonal populations. (E) Fraction of cells expressing Foxp3 for each CD4+ T cell clonal population. Clonal populations express Foxp3+ if at least one cell is Foxp3+. (F) Violin plot comparing the effector Treg geneset score of Foxp3+ cells from Foxp3 expressing CD4+ T cell clonal populations. (G-I) The CD8+ T cell clonal populations with cell members in both the tumour and spleen were assessed for differences in the differentiation of their cells across the two tissue compartments. (G) The mean exhaustion geneset score of cells from each clonal population plotted for each tissue compartment. Scores from the same clonal population are linked by a line. (H) Violin plot comparing the exhaustion geneset score of individual cells in G2M or S phase in each tissue compartment. (I) The mean exhaustion geneset score of cells from each clonal population plotted against the mean tissue resident memory score of cells. (J-K) The Treg clonal populations with cell members in both the tumour and spleen were assessed for differences in the differentiation of their cells across the two tissue compartments. (J) The mean effector Treg geneset score of cells from each clonal population plotted for each tissue compartment. Scores from the same clonal population are linked by a line. (K) Violin plot comparing the effector Treg geneset score of individual cells in G2M or S phase in each tissue compartment. Dots represent a single cell (F, H, K) and individual clonal populations (C, E, G, I, J). Adjusted R-squared and F test p values as shown for all clonal populations (I). Statistical testing via paired two-tailed students t-test and Kruskal-Wallis test (****, p <0.0001; ***, p <0.001; **, p <0.01; *, p<0.05; ns, p>0.05). (AU) arbitrary units.

We compared CD8+ T cell and Treg clonal population differentiation states across the spleen and the tumour. Advances in sequencing technologies have enabled the capture of clonally related cells across different tissues, but the paucity of antigen responding T cells in tissues such as the spleen, presents a significant sampling challenge. Using the AgRSR to fate-map and sort EYFP cells enables a 100-fold enrichment for these cells in the spleen, permitting a comprehensive comparison of clonally related T cells across tissues. In our dataset, all CD8+ T cell and Treg clonal populations contained cells expressing Nr4a1 (antigen signalling) and Ifng/Gzmb in the tumour (Figure S3A), confirming their relevance to the tumour response in silico. We computed the exhaustion levels of individual CD8+ T cells by scoring the expression of exhaustion associated genes(*34*) against a control group of genes(*35*). For CD8+ T cell clonal populations distributed across the spleen and the tumour, cells in the tumour, on average, had higher exhaustion levels than counterparts in the spleen (Figure 3G). Cells defined by their gene expression to be in a cycling phase, in the tumour, were more exhausted than counterparts in the spleen (Figure 3H). A group of clonal populations were prescribed lower exhaustion scores in the tumour (Figure 3H). These clonal populations expressed genes associated with a stem-like state(*36*) and did not expand as much as other clonal populations (Figure S3B). Across all CD8+ T cell clonal populations, the mean exhaustion score of cells in a clonal population strongly correlated with a clonal population’s tissue-residency score(*37*), a differentiation state associated with tissue confinement, even after overlapping genes from the genesets were removed (Figure 3I). We computed the effector Treg level using an effector Treg differentiation signature(*38*). In parallel to CD8+ T cell clonal populations, for Treg clonal populations distributed across the spleen and the tumour, cells in the tumour, on average, had higher effector Treg levels than counterparts in the spleen (Figure 3J). Cells in a cycling phase in the tumour were also in a higher effector Treg state than equivalent cells in the spleen (Figure 3K). These findings for CD8+ T cell and Treg clonal populations were reproduced when we utilised exhaustion and effector Treg genesets obtained in different immune settings(*39*–*44*), and replicated when we assessed the differentially expressed genes of splenic and tumour cells (Figures S3C, S3D and S3E). Taken together, these data indicate a spatial partitioning of tolerance between the effector site and the circulating immune system, universally adopted by T cells receiving systemic antigen signals.

Immune selection of cancer cells that have either deleted neoantigens(*45*, *46*) or by other mechanisms prevented MHC class I antigen presentation(*46*–*50*) at metastatic sites indicate that metastatic protection is T cell dependent. However, CD8+ T cells differentiate to an exhausted state after persistent antigen signalling – an epigenetically propagated, permanent state of tolerance(*51*, *52*). It is therefore unclear how antigen-signalled T cells constrain tolerance to the effector site and avoid compromising systemic protection.

We sought to address this question by tracking T cells receiving antigen signalling in the tumour. Recent works using the Kaede mouse have tracked T cells in the tumour through photoconversion(*53*). Conclusions for antigen responding T cells, however, have been limited by the mouse’s indiscriminate marking of tumour-antigen agnostic bystander cells, and its inability to track the progenies of labelled cells. We utilised the AgRSR mouse to enrich for T cells receiving antigen signalling in the tumour and tracked their clonal expansions. EYFP+ T cells from the tumour and spleen were collected 8 days after direct tumoral 4-hydroxytamoxifen injection, and cells from 3 mice were analysed by paired scRNA-seq and T-cell receptor (TCR) sequencing (Figure 4A). Fate-mapped CD8+ and CD4+ T cells produced expanded clonal populations (Figure 4B). In line with a previous study(*54*) finding limited activation of conventional CD4 T cell clones in the tumour, almost all the CD4+ T cell clonal populations persistently expressed Foxp3 and their Foxp3+ cells similarly upregulated genes associated with the effector Treg state (Figures 4C, 4D and 4E).

**Figure 4.**
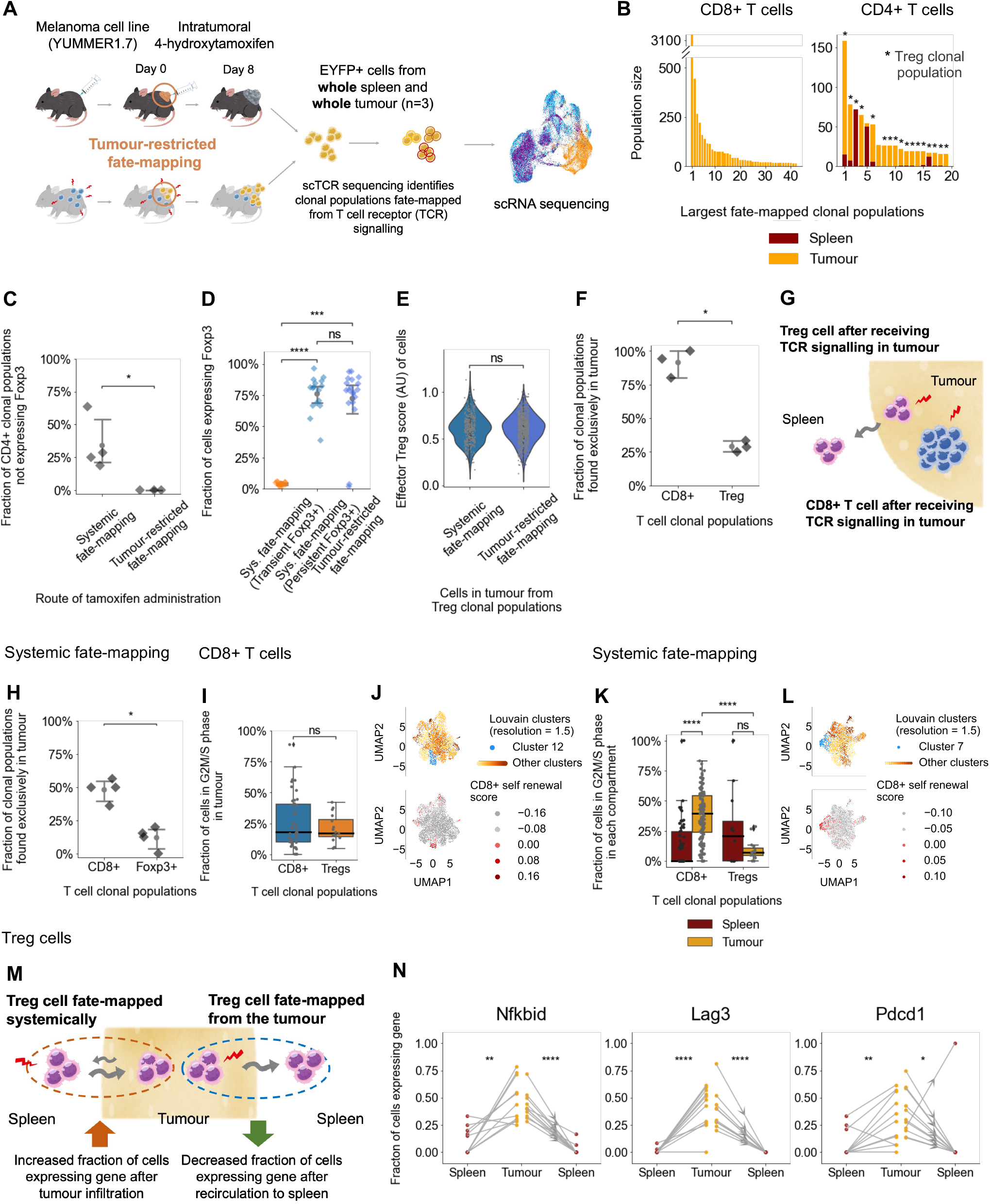
Distinct spatial CD8+ T and Treg cell behaviours localise tolerance to the tumour. (A) Schematic of experimental design; 8 days after intratumoral 4-hydroxytamoxifen injection EYFP+ T cells were sorted from the whole spleen and tumour of AgRSR-LSL-EYFP mice and subject to single-cell RNA and VDJ analysis (n=3). (B) Largest (population size greater than 15) fate-mapped CD8+ and CD4+ T cell clonal populations ranked by size and coloured by the numbers of cells within the tumour or spleen compartments. (C) Fraction of CD4+ T cell clonal populations not expressing Foxp3 in intraperitoneal (n=4) and intratumoral tamoxifen labelled mice (n=3). (D) Fraction of cells expressing Foxp3 for each CD4+ T cell clonal population. (E) Violin plot comparing the effector Treg geneset score of cells in the tumour of Treg clonal populations between intraperitoneal and intratumoral tamoxifen labelled mice. (F) Fraction of CD8+ T cell and Treg clonal populations, from intratumoral tamoxifen labelled mice, for which no cells were identified in the sorted EYFP+ cells of the spleen. (G) Schematic summarising the recirculation readiness of T cell clonal populations. (H) Fraction of CD8+ T cell and Treg clonal populations, from intraperitoneal tamoxifen labelled mice, for which no cells were identified in the sorted EYFP+ cells of the spleen. (I-L) Characteristics of CD8+ T cell clonal populations obtained from intratumoral (I-J) and intraperitoneal (K-L) tamoxifen labelled mice. (I) Boxplots showing the fraction of cells in the tumour in G2M or S phase. (J) UMAPs of tumoral CD8+ T cells coloured by Louvain clusters (Top) with highlighting of cluster highly expressing a self-renewal geneset (Bottom). (K) Boxplot showing the fraction of cells in G2M or S phase for CD8+ T cell and Treg clonal populations within the indicated tissue compartment. (K) UMAPs of tumoral CD8+ T cells coloured by Louvain clusters (Top) with highlighting of clusters highly expressing a self-renewal geneset (Bottom). (M) For each clonal population, and for each gene, we computed the fraction of cells expressing the gene in the spleen and the tumour. A group of genes was more frequently expressed in the tumour in both intratumoral and intraperitoneal tamoxifen labelled mice. (N) Three example genes. Clonal populations obtained from intratumoral tamoxifen labelled mice are shown with arrows. Dots represent a single cell (E), individual clonal populations (D, I, K, N) and individual mice (C, F, H). Statistical testing via paired two-tailed students t-test and Kruskal-Wallis test (****, p <0.0001; ***, p <0.001; **, p <0.01; *, p<0.05; ns, p>0.05). (AU) arbitrary units.

In this experiment, we sampled the whole spleen and tumour, enabling us to assess the spatial distribution of CD8+ T cell and Treg clonal populations without bias. Strikingly, while the majority of Treg clonal populations contained cells in the spleen, we captured only one CD8+ T cell from one CD8+ T cell clone outside of the tumour (Figure 4F). Extra-tumoral leakage of tamoxifen would have increased extratumoural EYFP+ CD8+ T cells suggesting that there was minimal leakage and strengthening a conclusion that trapped progeny results from intratumoural antigen-selection of CD8+ T cells. These results indicate that after antigen signalling in the tumour, Treg cells more readily recirculate back to the spleen and the circulatory immune system than CD8+ T cells (Figure 4G). We assessed whether these contrasting T cell spatial behaviours were evident after systemic fate-mapping. Half of the identified CD8+ T cell clonal populations had cellular membership confined to the tumour, whereas almost all Treg clonal populations contained cells in the spleen (Figure 4H). These results indicate that CD8+ T cells do not readily recirculate, but instead, accumulate in the tumour to form and maintain tumour-exclusive populations. Consistent with this idea, both fate-mapping strategies revealed CD8+ T cells to proliferate highly in the tumour, and to be clonally related to a tumour population of Tcf7 expressing stem-cell like cells(*36*) (Figures 4I, 4J, 4K and 4L). We compared the Tregs obtained from tumour-restricted fate-mapping with counterparts obtained from systemic fate-mapping (FigureS4A). In the spleen, Tregs from tumour-restricted fate-mapping expressed lower levels of genes associated with tumour homing (Ccr4, Cxcr3) (Figure S4B), consistent with tumour-restricted and systemic fatemapping strategies predominantly capturing recirculating and infiltrating Tregs respectively. We investigated how Treg states change after tumour infiltration and recirculation at a cellular level, by assessing the differentially expressed genes (Figure S4C) and at a clonal level, by comparing the changes in the fraction of cells expressing a particular gene for each clonal population, across the spleen and the tumour (Figure 4M). Unbiased assessment at the clonal level revealed a group of 346 genes expressed more frequently after tumour infiltration, and less frequently after recirculation. Gene ontology (GO) enrichment analysis associated a subset of these genes with TCR signalling (Nfkbid, Itk, Pde4d), and tolerance induction (Lag3, Pdcd1, Cd274) (Figures 4N, S4D and S4E). Together, these results indicate that after antigen signalling in the tumour, Tregs more readily recirculate than CD8+ T cells, but after recirculation and away from antigen, transition to a lower tolerance state.

To understand how the spatial partitioning of tolerance is temporally maintained, we tracked T cell clonal populations over time. Elegant studies based on adoptive transfer models have investigated the temporal characteristics of T cell responses in vivo, but difficulties in distinguishing separate clones, have prevented conclusions at a detailed temporal and spatial resolution. We systemically marked T cells receiving antigen signalling by intraperitoneal tamoxifen injection and sorted EYFP+ T cells from the tumour and spleen 18 days afterwards. Cells from 3 mice were analysed by paired scRNA-seq and T-cell receptor (TCR) sequencing, and these data were combined with previous analysis of mice at day 8 (Figure5A). For all T cells, most cells in the spleen remained non-expanded whereas most cells in the tumour were members of expanded clonal populations (Figure S5A). Again, we focused on the largest, expanded T cell clonal populations (Figure 5B). We identified the previous three CD4+ T cell clonal populations categories, as defined by Foxp3 expression (Figures 5C, S5B and S5C). Foxp3+ cells from populations that persistently expressed Foxp3 continued to differentially upregulate genes associated with effector Treg function, demonstrating their stable Treg lineage nature (Figure S5D). By day 18, the size of populations that transiently expressed Foxp3 was significantly larger in the spleen, and the size of populations that persistently expressed Foxp3 was significantly larger in the tumour (Figure S5E). Foxp3+ cells from populations that transiently expressed Foxp3 were no longer found in the tumour and were absent from all tissues in one mouse (Figures S5F and S5G). Both CD8+ T cell and Treg clonal populations maintained the tissue-dependent tolerance for antigen, as seen on day 8 (Figures 5D, 5E, 5F, 5G, 5H, S6A and S6B). We analysed changes in the CD8+ T cell and Treg clonal populations between days 8 and 18. In CD8+ T cell clonal populations, the fraction of cycling cells decreased, with almost no clonal populations containing cycling cells in the spleen by day 18 (Figure 5I). A lower fraction of cells from CD8+ T cell clonal populations were also found in the spleen (Figure 5J). Compared to day 8, day 18 populations expressed higher levels of genes associated with exhaustion (Figures 5K and S6C), corroborating previous reports linking increasing exhaustion with persistent antigen exposure(*55*). The increase in exhaustion was primarily driven by T cells that were members of clonal populations found exclusively in the tumour (Figure S6D), suggesting tumour residency, and lack of supply from the periphery drive populations to higher levels of exhaustion. Since peripheral cells are associated with non-exhausted, cytotoxic function(*56*), we hypothesised that reduction of splenic population would lead to loss of the most cytotoxic cells. Although some clones within the tumour appeared cytotoxic and functional on day 8, these did not exist at day 18 (Figure 5L). In contrast, in Treg clonal populations, neither the fraction of cycling cells, nor the clonal population’s spatial distribution, nor the effector Treg score changed significantly between days 8 and 18 (Figures 5M, 5N, 5O and S6E).

**Figure 5.**
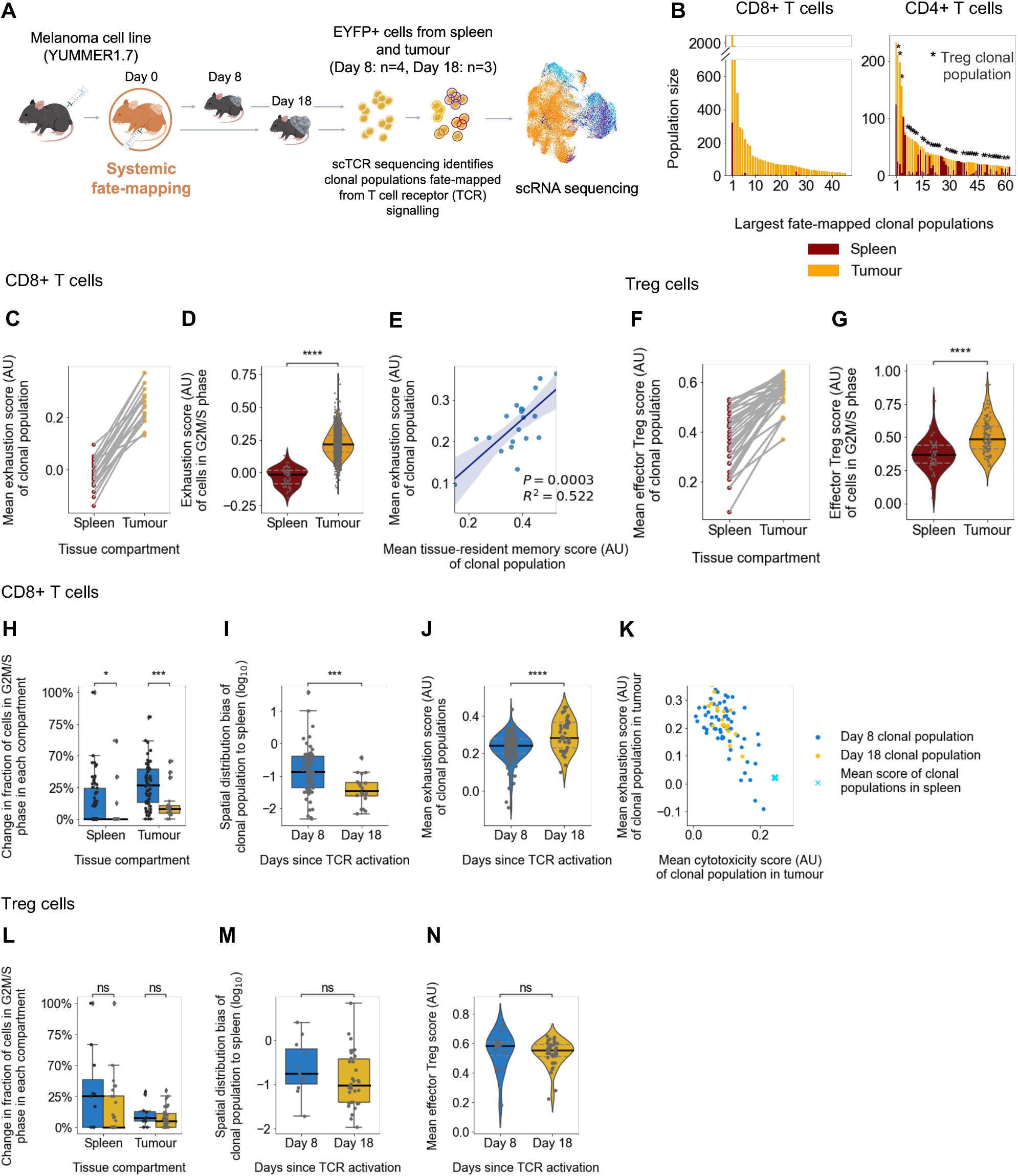
Distinct temporal CD8+ T and Treg cell behaviours localise tolerance to the tumour. (A) Schematic of experimental design; 8 and 18 days after intraperitoneal tamoxifen injection EYFP+ T cells were sorted from the spleen and tumour from AgRSR-EYFP-LSL mice and subject to single-cell RNA and VDJ analysis (n=4 and n=3 respectively). (B-G) Analysing the T cell clonal populations collected at day 18. (B) Largest (population size greater than 15) fate-mapped CD8+ and CD4+ T cell clonal populations, ranked by size and coloured by the numbers of cells within the tumour or spleen compartments. (C-G) The CD8+ T cell (C-E) and Treg (F-G) clonal populations with cell members in both the tumour and spleen were assessed for differences in the differentiation of their cells between different tissues. (C) The mean exhaustion geneset score of cells from each CD8+ T cell clonal population plotted for each tissue compartment. Scores from the same clonal population are linked by a line. (D) Violin plot comparing the exhaustion geneset score of individual cells in G2M or S phase in each tissue compartment. (E) The mean exhaustion geneset score of cells from each clonal population plotted against the mean tissue resident memory score of cells. (F) The mean effector Treg geneset score of cells from each clonal population plotted for each tissue compartment. Scores from the same clonal population are linked by a line. (G) Violin plot comparing the effector Treg geneset score of individual cells in G2M or S phase in each tissue compartment. (H-N) Tracking how the spatial characteristics of CD8+ T cell (H-K) and Treg (L-N) clonal populations change over days 8 and 18. (H) Boxplot comparing the fraction of cells in G2M or S phase in the spleen and tumour for CD8+ T cell clonal populations at days 8 and 18. (I) Boxplot comparing the spatial distribution bias of CD8+ T cell clonal populations to the spleen at days 8 and 18. The spatial distribution bias was calculated by dividing the spleen count frequency by the tumour count frequency for each clonal populations. A high spatial distribution bias indicates a clonal population’s bias to the spleen, a low number indicates bias to the tumour. (J) Violin plot comparing the exhaustion geneset score of CD8+ T cell clonal populations collected at days 8 and 18. (K) Transcriptional assessment of the mean exhaustion and cytotoxic geneset score of cells from each CD8+ T cell clonal population, restricted to analysis of cells from the tumour (Small dots) or the spleen (Cross). The mean exhaustion geneset score is coloured by time since tamoxifen. (L-N) Treg clone population equivalent of (H) and (I) respectively. (N) Violin plot comparing the effector Treg geneset score of Treg clonal populations collected at days 8 and 18. Dots represent a single cell (D, G) and individual clonal populations (C, E, F, H, I, J, K, L, M, N). Adjusted R-squared and F test p values as shown for all clonal populations (E). Statistical testing via paired twotailed students t-test and Kruskal-Wallis test (****, p <0.0001; ***, p <0.001; **, p <0.01; *, p<0.05; ns, p>0.05). (AU) arbitrary units.

## Discussion

In this study, we develop and validate a novel fate-mapping mouse to track lymphocytes based on induced antigen signals. Our system overcomes inadequacies of previous technologies by enabling the marking of lymphocyte clones responding to all types of antigens, including undefined and endogenous antigen, while also distinguishing the variability of antigen signalling due to accessory signalling. We successfully validate the system’s specificity for exclusively marking lymphocytes (and their progenies) that have received antigen signalling *in vivo*.

Using the AgRSR system to study the tolerance of tumours, we report how contrasting T cell spatial and temporal behaviours combine to constrain T cell tolerance to a particular anatomical niche. Tregs recirculate and ‘resupply’ their peripheral population, differentiating to a reversible tolerant state within the tumour, while CD8+ T cells less readily recirculate, ‘lose’ their peripheral population, and differentiate to an irreversible, increasingly exhaustive, tolerant state in the tumour. These findings suggest interventions that promote CD8+ T cell recirculation and prevent Treg recirculation could synergise with immunotherapy in the treatment of cancers and other diseases caused by antigen-persistence by breaching the tissue-confined tolerance.

More broadly, these T cell behaviours provide a mechanism by which immunity can tolerate a particular anatomical niche while maintaining systemic clonal protection. The antigen-receptor of T cell clones, the TCR, relies on antigenic-structure to determine clone selection. It cannot, by itself, evaluate the pathogenicity of the antigenic source nor the load and distribution of the antigen. Sustained work over the last three decades has demonstrated how the former constraint is overcome by innate immune recognition signals, but no mechanism has been proposed to address the latter constraint. Our work presents spatiotemporal behaviours of T cell clones that enable tissue-specific responses to locally high and persistent antigen signalling. In the face of ineliminable pathogen, these clonal behaviours may have evolved to prevent tissue-destructive responses at one site while enabling ongoing protection at other sites.

We stress that these novel findings were only detected by the ability of the AgRSR mouse to mark clones that had received antigen signalling at discrete times and locations. Alongside capacity for fate-mapping CD8+ and CD4+ T cells, this system could track (self-) antigen-stimulated B lymphocytes. We envision the system to probe fundamental questions of immunity that could have profound impact on our understanding of health.

## Acknowledgements

This work has been funded by the Medical Research Council (MC_UU_0025/12) and Medical Research Foundation (MRF-057-0002-RG-THAV-C0798) awards to JEDT. DIJ reports grants from Cancer Research UK, supporting his laboratory work. From April 2019, the core grant to the CI was C9545/A29580. The Li Ka Shing Centre where components of this work was performed was generously funded by CK Hutchison Holdings Limited, the University of Cambridge, Cancer Research UK, The Atlantic Philanthropies and others. PAL and KGCS additionally supported by EU H2020 research and innovation programme under grant agreement 668036 (RELENT). MT is supported by the Masason Foundation and the Gates Cambridge Trust. We thank Melania Barile and Xianon Wang from the Gottgens group for their insights into single cell analysis. We thank the Core Facilities at the Cancer Research UK Cambridge Institute (Li Ka Shing Centre), including the Biological Resources Unit, Research Instrumentation & Cell Services, Light Microscopy and Flow Cytometry for technical provision.

## Author contributions

J. E.D.T. designed and developed the AgRSR mouse. T.Y.S conducted experiments with the assistance of K. K., M.R., T.M., M.S., and P.C. M.T. conducted the single-cell analysis with the assistance of K.W. and Z.H. M.T., P.L.. J.E.D.T. supervised the work with the assistance of D.N., T.H., P.A.L., P.L., K.O., D.A., P.A.L., K.G.C.S., D.I.J. and M.A.C. All authors contributed to the analysis of the presented results. J.E.D.T and M.T. wrote the paper with input from all other authors. J.E.D.T. conceived the research programme.

## Corresponding Authors

Correspondence and requests for materials should be addressed to J.E.D.T.

## Methods

### BAC Clone Modification and Purification

A BAC clone containing the *Nur77* gene (BACPAC resources) was modified by introducing Katushka E2A linked CreERT2-SV40 polyadenylation signal into the start ATG of the *Nur77* gene by homologous recombination(*57*). BAC DNA was purified from 200ml bacterial cultures by alkaline lysis (Qiagen buffers), and circular DNA was separated by CsCl ultracentrifugation. Briefly, 4.04g of CsCl were added to 4ml resuspended DNA and CsCl dissolved at 40°C. 25 μl 10mg/ml EtBr and 75μl water was added. Samples were spun in a bench tube centrifuge at 3000 rpm for 15 minutes to remove remaining proteins. The DNA CsCl solution was spun at 70,000g for 6 hours. EtBr was removed by n-butanol extraction and the DNA precipitated. Successful recombination was confirmed by PCR. The DNA was spot dialyzed on Millipore VSWPO2500 filters into polyamine buffer (10mM Tris-Cl, pH 7.5, 0.1mM EDTA, 100mM NaCl, 30μM spermine, and 70μM spermidine).

### Experimental Mice

AgRSR mice were generated via pronuclear injection of the modified BAC DNA into 0.5 d fertilized ova of C57BL/6 donors. Founder lines were assessed for transgene expression and the line with the highest expression was crossed with the ROSA26-LSL-EYFP mice (gift from Prof. Doug Winton, CRUK-CI, Cambridge). The AgRSR mice and B2m^-/-^ (Jax, 002087), RAG2^-/-^ (Jax, 008449) animals were maintained in CRUK Cambridge Institute Biological Resources Unit and University of Cambridge Central Biomedical Service, under specific-pathogen-free conditions. Genotyping was performed by Transnetyx using established probes. All animal experiments were conducted when mice were between 8-12 weeks of age and were conducted in accordance with Home Office guidelines. xx non-transgenic mouse were used as controls to determine Katushka labelling specificity.

### Listeria, Vaccine and Tumour Challenges

For listeria infection, mice were infected with 1500cfu of *Listeria moncytogenes* in experiments indicated in the text. For vaccination, mice were immunised by intraperitoneal injection of 50μg SIINFEKL peptide, 10μg anti-CD40 (Bio X cell) and 10μg Poly:IC (InvivoGen). For the generation of murine tumours, cells of the cultured YUMMER1.7 cell line(*31*) (a gift from M. Bosenberg) were detached with 0.5% trypsin-EDTA (Gibco) for 3 minutes, quenched with complete media, and washed in PBS three times. Single-cell suspensions of 1 million cells were subcutaneously injected into the right flank of each mouse. In vivo tumor volumes were monitored by (Width x Depth x Length)/2 using a calliper.

### Tamoxifen Administration

Tamoxifen (20mg/ml) was first prepared in EtOH (5-10% v/v) and sunflower oil (90-95% v/v) before dissolving in a 37°C water bath under sonication (35kHz) for 15 minutes. Tamoxifen (2mg) was administered by intraperitoneal injection 24 hours after the *Listeria* challenges, at 0 and 12 hours after the vaccine challenges, 5 days before tumour implantation in the tamoxifen pre-tumour challenge or at day 7 after subcutaneous tumour implantation in all other tumour experiments. For fate-mapping TCR activation within the tumours, 10μl of 4-hydroxytamoxifen (4OHT) at (39mg/ml) was injected into tumours at day 10 after subcutaneous tumour injection. Mice in which 4OHT extratumorally leaked were excluded.

### Generation of Single-Cell Suspensions from Tissues

Inguinal lymph nodes and spleens were homogenized in PBS/ 0.1% FCS/ 2mM EDTA and filtered (70μm or 100μm filters). RBC lysis buffer (NH4Cl/ NaHCO3/ EDTA) was used to lyse splenic erythrocytes. Tumours were cut into pieces by a scissor and digested using the Miltenyi Tumour Dissociation Kit, according to the manufacturer instructions, before filtering (100μm then 70μm) to generate a single cell suspension. For subsequent culture, T cells were selected from single-cell spleen suspension using a magnetic cell separation (LS column, Miltneyi).

### In vitro cell culture

Purified T cells from the spleen were cultured in complete RPMI (10% FBS, 55 mM 2-Mercapthoethanol). T cell media was supplemented with IL-2 (20ng/ml), IL-7 (2ng/ml) and the SIINFEKL peptide and its variants at (5ug/ml), and incubated at 37°C and 5% CO2 for 24 hours.

### Flow Cytometry

Surface antibody staining was conducted in PBS containing 0.1% FCS and 2mM EDTA for 30 minutes at 4°C to cells that had been pre-incubated with Fc block and LIVE/DEAD Blue (InVitrogen) staining. Samples were fixed by incubating in buffer containing 1% formaldehyde/0.02% sodium azide/glucose/PBS for 10 minutes at room temperature. Data was acquired on a LSR Fortessa (BD Bioscience) flow cytometer and further analysed by Flowjo v10 (Treestar).

### Preparation of EYFP+ Cells for scRNAseq

Cell suspensions from tumour, spleen, and inguinal lymph nodes were generated independently. Samples were stained as for FACS analysis with CD45-APC, CD11b-PeCy7, CD19-PeCy7, F4/80-PeCy7 antibodies for 30 minutes on ice. Live immune cells were sorted using a FACS Aria instrument (BD Bioscience) with a 100μm nozzle. The sorted cells were collected in PBS containing 10%FCS and 2mM EDTA in cold and spun down at 300g 7minutes 4°C before resuspending in PBS to achieve a final concentration of 10-20,000 cells/32μl. Totalseq C Hashtag antibodies (1, 2, 3) were used to barcode inguinal lymph nodes (tumour-draining and contralateral) and spleen samples to enable pooled library preparation. Hashtags were added to individual tissues at 0.1mg/ml; cells were then washed twice and sorted. Hash-tagged samples were pooled just prior to scRNAseq to minimise hashtag carryover. In experiments fate-mapping TCR activation with, intratumoral 4OHT, CD4+ and CD8+ lymphocytes were enriched from tumour and spleen single cell suspensions by magnetic bead purification using Miltenyi beads. Cells were then stained with antibodies against CD11b, CD19, F4/80 and CD45.2 according to our FACS staining protocol above, with the addition of TotalseqC Hashtag antibodies against CD45 and H2Kb (1:500). Samples were then washed twice and EYFP+ T cells (lineage negative cells) were sorted. Hash-tagged samples were pooled just prior to scRNAseq to minimise hashtag carryover.

### Single Cell Library Preparation and Sequencing

Single-cell RNA-seq libraries were prepared using Chromium Single Cell and V(D)J Enrichment Kits following the Single-Cell V(D)J Reagent Kits User Guide (Manual Part CG000086 Rev H, I, J, K, L, M; 10X Genomics). The data from the experiment using intraperitoneal tamoxifen were generated using chemistry (5’ v 1) before the introduction of dual-indexing strategy; the data for the intratumoural 4OHT experiment was generated using chemistry (5’ v 2). Sorted samples were resuspended in PBS-0.04% BSA and loaded into Chromium microfluidic chips to generate single-cell gel-bead emulsions using the Chromium controller (10X Genomics). RNA from the barcoded cells for each sample was reverse transcribed in a C1000 Touch Thermal cycler (Bio-Rad), and libraries generated according to the manufacturer’s protocol with no modifications (14 cycles used for cDNA amplification). Library quality was confirmed with Agilent TapeStation 4200 (High Sensitivity D1000 ScreenTape to evaluate library sizes) and BMG LABTECH Clariostar Monochromator Microplate Reader (Invitrogen Quant-iT dsDNA Assay Kit; high sensitivity to evaluate dsDNA quantity). Samples were sequenced on an Illumina HiSeq 4000 as 2 × 150 paired-end reads, one full lane per pool (before analysis, gene expression data were trimmed to 26 bp, read 1; 8 bp, i7 index; and 98 bp, read 2) or run on Illumina NovaSeq6000 with the same parameters (PE150, gene expression libraries trimmed to 28:8:0:98).

### Software versions

Data was analysed using R version 4.0.3 and R packages (Seurat 4.0.4, SeuratData 0.2.1, SeuratDisk 0.0.0, scRepetoire 1.0.0 and monocle3 0.2.3), Python version 3.8.6 and Python packages (jupyterlab 2.2.9, numpy 1.19.4, pandas 1.1.5, scipy 1.6.0, scanpy 1.6.0, anndata 0.7.5, rpy2 3.3.6, anndata2ri 1.0.5.dev2+ea266ab and scikit-learn 0.24.1). Figures were produced using seaborn 0.11.0, matplotlib 3.5.1 in Python, EnhancedVolcano 1.11.3 in R, Prism 10 and Adobe Illustrator 26.2.1.

### Preprocessing of scRNA-seq data

10x Genomics gene expression raw sequencing data were processed using CellRanger software v.3.0, and the VDJ TCR alpha and beta chains were processed using CellRanger VDJ v.3.1.0, both following the CellRanger pipeline. Sequencing reads were aligned to the mouse reference genome mm10 (Ensembl 93) provided by CellRanger. To prevent the variable TCR genes from contributing to downstream analysis, genes containing Trav and Trbv from their gene name were removed from the gene-expression matrix. The resulting count matrices and VDJ sequences were further processed using Seurat v3.2.3(*58*) and scRepetoire v1.0.0(*59*). In brief, samples were demultiplexed with the aid of the HTODemux function, matched to their VDJ sequences with the combineTCR and combineExpression functions, filtered (to cells that had full TCR alpha and beta nucleotide sequences, genes in the 200-5000 range, and less than 10 percent mitochondrial genes), pre-processed by regressing out cell cycle scores and applying the SCTransform function for each sample, and integrated with the FindIntegrationAnchors and IntegrateData functions(*60*). The days 8 and 18 intraperitoneal injection experiment integrated data from 4 (38356 cells after integration) and 3 (26697 cells after integration) mice respectively, and the day 8 intratumoral injection experiment integrated data from 3 mice (10793 cells after integration).

### Clonal Populations

We defined cells with identical TCR alpha and beta nucleotide sequences as cells belonging to the same clonal population. If the raw expression count of a gene (e.g. CD8a, CD4, Foxp3) was non-zero, cells were defined to be expressing that gene. Clonal population frequencies in each tissue compartment were calculated by dividing the number of cells by the total number of T cells captured in the respective tissue compartment for each sample. The ‘spatial distribution bias of clonal population to the spleen’ was calculated by dividing the spleen population frequency by the tumour population frequency for each clonal population.

### Geneset Scores

The genesets for cell cycle status were taken from Tirosh et al., 2016(*35*) and the CellCycleScoring module was used to assign cells either a G1 or G2M/S phase (proliferating) status in Seurat. Genesets from Li et al., (2018) (*34*), Bending et al., (2018)(*38*) Mackay et al., (2013)(*37*) and Yost et. al (2019) (*61*) (based on Im et al., (2017)(*36*) were used to score cells for exhaustion, effector Treg differentiation, T cell resident memory and Tcf7 self-renewal capacity respectively. Additional genesets from Lucca et al., (2021)(*39*), Wherry et al., (2007)(*41*) and Beltra et al., (2020)(*62*) (based on Bengsch et al., (2018)(*40*)) were used to corroborate exhaustion scores, and genesets from Perry et al., (2022)(*42*), Dias et al., (2017)(*43*) and Ouyang et al., (2012)(*44*). Mouse ortholog genes (from http://genomespot.blogspot.com/2016/12/msigdb-gene-sets-for-mouse.html, Ensembl Biomart version 87) were used when genesets were derived from human data. Geneset scores were calculated by normalising and logarithmising the raw gene counts obtained from the CellRanger output and applying the score_genes function from Scanpy on the relevant dataset, as shown in Tirosh et al., 2016 (*35*). Scores were produced for each cell, and these were then averaged for cells in a clonal population (or other subsets of cells as indicated in the text) to calculate scores for each clonal population.

### Clustering / UMAPs of Data

For both day 8 intraperitoneal and day 8 intratumoral injection experiments, CD8+ T cells from the tumour were subset from the main dataset, and nearest neighbourhood graphs (size of local neighborhood=15), Prinicipal Component Analysis (PCA) coordinates (number of dimensions = 50) and Uniform Manifold Approximation and Projection (UMAP) coordinates were generated using functions from Scanpy. Louvain clustering (*63*) was applied to each dataset at resolutions of 1.0 and 1.5 respectively to perform unsupervised clustering of the data.

### Statistical Analysis

Statistical analyses on the flow analysis data were performed using two-tailed Student’s t-tests unless otherwise indicated. For the single cell data, regression lines and corresponding confidence intervals were calculated using the regplot parameter from Seaborn. Differential gene expression was computed using the FindMarkers function (wilcox test, logfc.threshold set to 0.25) from Seurat. Adjusted R-squared score and F test p values were computed using linear least-squares regression, and paired comparisons were assessed for statistical significance using Kruskal-Wallis with Bonferroni correction in Scipy.

